# SSI: A Statistical Sensitivity Index for Chemical Reaction Networks in cancer

**DOI:** 10.1101/2023.01.12.523784

**Authors:** Giorgia Biddau, Giacomo Caviglia, Michele Piana, Sara Sommariva

## Abstract

**Summary:** At the cellular level, cancer is triggered by mutations of the proteins involved in signalling networks made of hundreds of reacting species. The corresponding mathematical model consists of a large system of non-linear Ordinary Differential Equations for the unknown proteins concentrations depending on a consistently large number of kinetic parameters and initial concentrations. For this model, the present paper considers the problem of assessing the impact of each parameter and initial concentration on the system’s output. More specifically, we introduced a statistical sensitivity index whose values can be easily computed by means of principal component analysis, and which leads to the partition of the parameters’ and initial concentrations’ sets into sensible and non-sensible families. This approach allows the identification of those kinetic parameters and initial concentrations that mostly impact the mutation-driven modification of the proteomic profile at equilibrium, and of those pathways in the network that are mostly affected by the presence of mutations in the cancer cell.

## 1. Introduction

The mathematical modeling of chemical reactions networks (CRNs) may involve a large number of differential equations and parameters (Zi, 2011). By instance, a model recently proposed for the colon-rectal cancer (CRC) incorporates the available biological knowledge into a system of 419 proteins interacting through 850 chemical reactions with as many rate constants (Tortolina *and others*, 2015; Santra, 2018; Sommariva and others, 2021*a*). In this model, under the assumption that the chemical reactions rely on mass action kinetics, the network accordingly corresponds to a non linear system of 419 ordinary differential equations (ODEs), involving 419 protein concentrations as unknowns, and 850 rate constants as equations’ coefficients (Sommariva *and others*, 2021*a*). Following an assignment of 419 initial conditions, the solution of the system of ODEs provides the simulated time course of the concentrations, and the (possibly allowed) asymptotic stationary state, to be identified with an equilibrium state. Numerical simulations may be used to predict changes in the model response resulting from varied external signals or internal conditions (Zi, 2011), to put into evidence feedback effects (Santra, 2018; Sommariva *and others*, 2021*b*), and to determine the impact of the introduction of targeted drugs (Santra, 2018; Krishnan *and others*, 2020; Sommariva *and others*, 2021*b*).

It is well known that the results obtained from simulations depend on the values assigned to the parameters and to the initial conditions, which, in the case of the CRC CRN in the G1-S phase (Bartek and Lukas, 2001), were fixed on the basis of literature data (Tortolina *and others*, 2015). This fact opens a crucial issue about how to quantitatively assess the impact of the parameters’ uncertainty on the CRC CRN equilibrium state. The present paper is devoted to a general analysis of this problem and illustrates a novel statistical tool for its solution arising from sensitivity analysis. In fact, our approach is specifically tailored to static conditions, which provides notable formal and computational advantages in contrast with the complexity of global dynamical approaches (Saltelli *and others*, 2000, 2005; Quaiser and Mönnigmann, 2009; Saltelli *and others*, 2010; Zi, 2011). Specifically, this analysis relies on two steps: an application of the implicit function theorem, whereby equilibrium states are obtained as functions of the parameters; and a principal component analysis (PCA), which leads to the distinction between sensible and non sensible rate constants and initial conditions via the definition of a statistical sensitivity index (SSI) weighing the relevance of either a specific constant or of the corresponding reaction. We point out that this second PCA-based step is inspired by an approach utilized by Liu *and others* (2005). However, differently than our case, that sensitivity index was introduced under dynamic conditions and, further, it suffered an intrinsic eigenvector sign ambiguity. The application of our analysis to the CRC CRN shows that SSI provides a reliable measure of the mutations’ impact on the protein expression, and allows the identification of the reactions that are mostly triggered by those mutations.

The plan of the paper is as follows. Section 2 setups the formalism of the mathematical description of a CRN and introduces SSI. Section 3 shows some results concerned with the CRC CRN. Our conclusions are offered in Section 4.

## 2. Methods

### 2.1 Equilibrium point for CRNs

We consider a CRN comprising *r* reactions involving *n* well-mixed chemical species and assume that the law of mass action kinetics holds. In the following we shall denote with *x*_*i*_(*t*), *i* = 1, …, *n*, the molar concentration (nM) at time *t* of the *i* −th species *A*_*i*_, and with *k*_*j*_, *j* = 1, …, *r*, the rate constant corresponding to the *j*−th reaction. The dynamic of the species concentration can be described by a system of *n* ordinary differential equations (ODEs) of the form

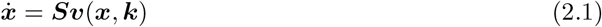

where 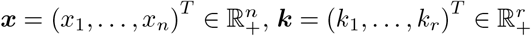 is the vector of rate constants, ***S*** ∈ ℤ^*n×r*^ is the stoichiometric matrix, 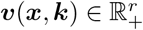 is the vector of reaction fluxes defined through the law of mass action, and the superimposed dot denotes the time derivative (Feinberg, 1987; Chellaboina *and others*, 2009; Yu and Craciun, 2018).

We further assume the CRN to be weakly elemented (Sommariva *and others*, 2021*a*), that is *p* = *n* − rank(***S***) conservation laws exist described by the linear system

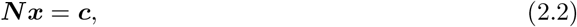

where 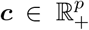 is the vector of conservations laws’ constants, and the matrix ***N*** ∈ ℕ^*p×n*^ is determined by computing the kernel of ***S***^*T*^ and is assumed to contain a minor equal to the identity matrix of size *p*. Therefore, up to a change in the order of the components of ***x***, we can consider the decompositions

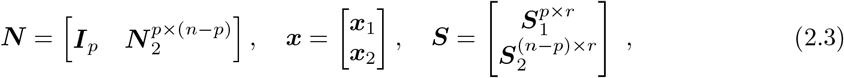

where ***x***_1_ is a *p*-vector of the so called elemental species (Shinar *and others*, 2009; Sommariva *and others*, 2021*a*) and ***S***_2_ ∈ ℝ^(*n*−*p*)*×r*^ has rank (*n* − *p*) and is the submatrix of ***S*** corresponding to the non-elemental species.

It follows from the definition of conservation laws that ***NS*** = 0, whence ***S***_1_ = −***N*** _2_***S***_2_. It can be shown (Chellaboina *and others*, 2009; Berra *and others*, 2022) that the equilibrium (stationary) solution of the Cauchy problem is determined by the algebraic system

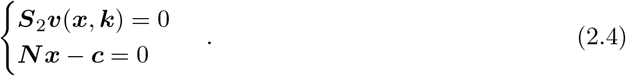

We observe that seeking the solution of this algebraic system is equivalent to the root-finding problem

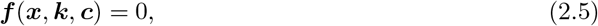

where *f* : ℝ^*n*^ × ℝ^*r*^ × ℝ^*p* →^ ℝ^*n*^ is a continuously differentiable function defined as

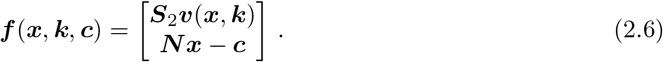

In the case of the CRC CRN, to any fixed ***k*** and ***c*** there corresponds a unique equilibrium/stationary state ***x***^*e*^, which may be determined by either a dynamic simulation (Sommariva *and others*, 2021*a*) or the solution of (2.4) (Berra *and others*, 2022). Moreover, since each reaction involves up to two reactants, from the law of mass action it follows that the components of ***f*** are polynomials of degree two or lower.

### 2.2 Analytic computation of the local sensitivity matrix

Local sensitivity analysis consists in determining how small variations of a parameter input influence the output of a certain model (Goulet, 2016; Varma and Morbidelli, 1999). In this particular case, we are interested in studying the local sensitivity of an equilibrium/stationary state ***x***^*e*^ with respect to the parameters ***k*** and ***c*** by constructing the local sensitivity matrix. Given a weakly elemented CRN, we denote with ***k***^0^ and ***c***^0^ the vectors whose components are the values of the network rate constants and conservation laws’ constants, respectively, when the network’s equilibrium is a given point ***x***^*e*^. Further, we define the vector ***h***^0^ = (***k***^0^, ***c***^0^) ∈ ℝ^*r*+*p*^, and we assume that

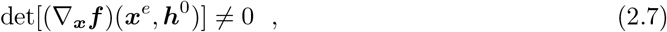

where henceforth, given a multivariate, vector-valued, continuously differentiable function ***g***, ∇_***z***_***g*** will denote the Jacobian matrix of ***g*** with respect to the variable(s) ***z***. On account of the implicit function theorem (Krantz and Parks, 2002), equations (2.4) and (2.7) imply that there exists one and only one ***x*** = ***x***(***h***) such that

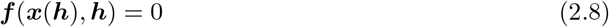

for ***h*** = (***k, c***) in a neighborhood of ***h***^0^, and thus ***x***(***h***) is the equilibrium point of the CRN defined by the set of parameters ***h***. Hence, the local sensitivity matrix is defined as

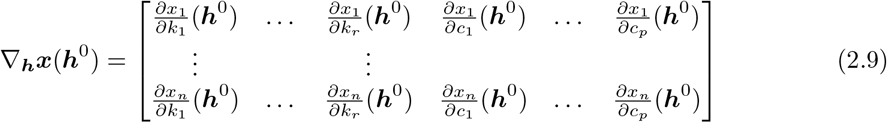

and can be computed by using

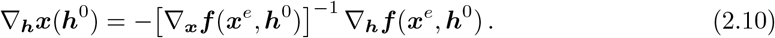

Further, the definition (2.6) of ***f*** implies that

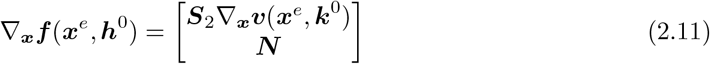

and

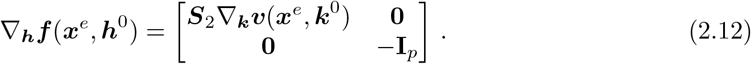

The submatrix of (2.9) formed by the first *r* columns is the sensitivity matrix with respect to the rate constants, while the submatrix formed by the last *p* columns is the sensitivity matrix with respect to the conservation laws’ constants.

### 2.3 Statistical Sensitivity Index

The first order Taylor expansion for the implicit function ***x*** = ***x***(***h***) leads to

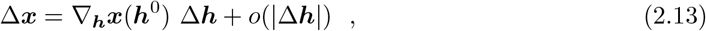

where

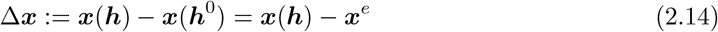

and

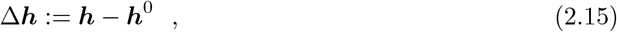

while the *n*×(*r*+*p*) matrix ∇_***h***_***x***(***h***^0^) is computed as in equation (2.10) and describes the variations of equilibrium points in terms of the variations of the set ***h*** of parameters.

By using (2.13) at first order and by applying some straightforward algebraic computations we obtained

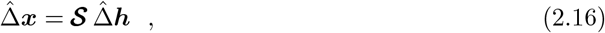

with

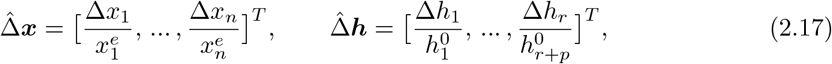

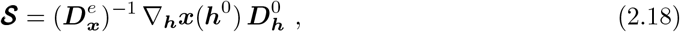

and

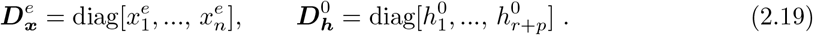

Following Liu *and others* (2005), we found that the function

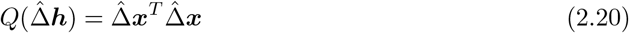

represents a measure of the relative deviation of concentrations resulting from a relative change in ***h*** at equilibrium. Clearly, *Q* is a positive semi-definite quadratic form, so that

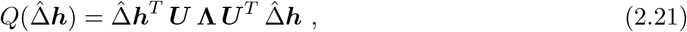

where the matrices **Λ** = diag[*λ*_1_, …, *λ*_*r*+*p*_], with *λ*_1_ ⩾ *λ*_2_ ⩾… ⩾ *λ*_*r*+*p*_ ⩾ 0, and ***U*** = [***u***_1_, …, ***u***_*r*+*p*_] are formed by the eigenvalues and the corresponding normalized eigenvectors of **𝒮** ^*T*^ **𝒮**, respectively. Equation (2.21) allows weighing the relevance of the *j*-th parameter, corresponding to the fractional change 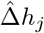. Indeed, when 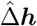 has 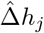 as unique non-vanishing component, (2.21) leads to

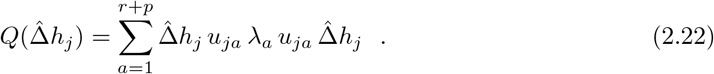

This equation inspires the definition of the *j*-th statistical sensitivity index (SSI)

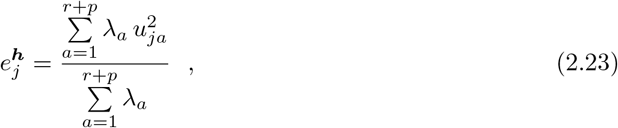

which is the average value of the squared components of the eigenvectors of **𝒮** ^*T*^ **𝒮** weighed by the corresponding eigenvalues, and whose values range in [0, 1]. Getting back to the original rate constant vector ***k*** and conservation laws’ vector ***c***, (2.23) naturally implies that 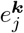 provides a synthetic and compact estimate of the weight of reaction *j* in the whole CRN, while 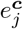 refers to the overall sensitivity of the network to the *j*-th component of vector **c**. For illustrative purpose, the entire workflow for computing the SSI values is summarized in Figure 1.

**Fig. 1.**
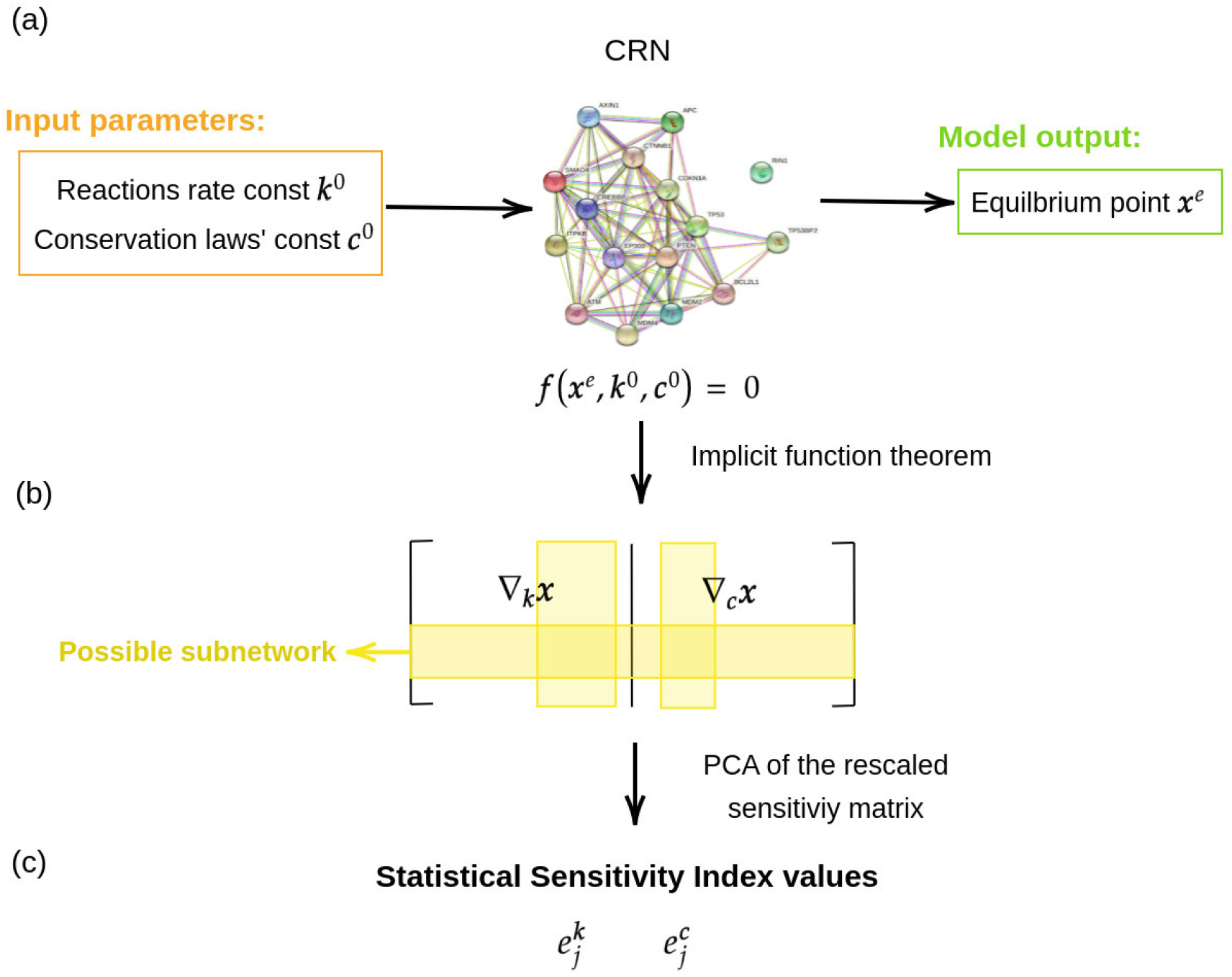
Schematic representation of the workflow for computing the SSI: (a) sensitivity analysis allows the quantification of how small changes in the input parameters, namely the rate constants ***k***^0^ and the conservation laws’ constants ***c***^0^, impact the protein concentrations ***x***^*e*^ at equilibrium; (b) by applying the implicit function theorem, the sensitivity matrix ∇_***h***_***x*** = [∇_***k***_***x***, ∇_***c***_***x***] can be analytically computed; (c) SSI values are then determined by performing a PCA of the whole rescaled sensitivity matrix or of a proper submatrix (yellow bands) representing a given subnetwork.

#### 2.3.1 Sensitivity analysis for a subnetwork

As schematically illustrated in Figure 1, SSI can be used also for the sensitivity analysis of subnetworks of a CRN. Indeed, let 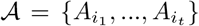, *t* < *n*, be the set of chemical species involved in a specific subnetwork. Then, the submatrix 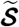 of **S**, formed by the *t* rows referred to 𝒜, is the searched sensitivity matrix. From this, the diagonalization of 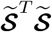 provides the SSI values referred to 𝒜. Specifically, such SSI values allow discriminating the parameters of the network whose variation mainly affects the values of the concentrations at equilibrium for the subset 𝒜 of chemical species.

*Remark* : Let us consider the case where t=1, i.e. *A* = {*A*_*i*_}, and let **s** be the *i*-th row of **𝒮**.

Given the diagonalization **s**^*T*^ **s** = ***U* Λ*U*** ^*T*^, then

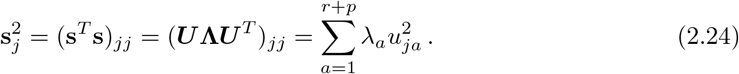

Hence, in this case, the statistical sensitivity indices can be computed as

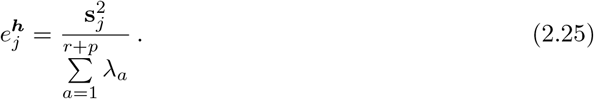

We notice that, since the denominator of equation (2.25) does not depend on *j*, the parameter rankings determined by SSI and by the square of the components of **s** coincide. This is coherent with the fact that the components of **s** are the elements of the rescaled sensitivity matrix **𝒮** describing the relative sensitivity of the equilibrium concentrations of the species 𝒜 with respect to the parameters of the network.

#### 2.3.2 Sensitivity analysis of a mutated CRN

The mathematical description of a pathological CRN can be obtained by implementing the presence of mutations in the Cauchy problem associated to the healthy CRN. When the Loss of Function (LoF) mutation of an elemental species occurs (Sommariva *and others*, 2021*b*), the corresponding conservation law is null. It follows that all the species involved in the conservation law are null during time. Let us call 𝒜 this set of species and ℬ the set of reactions where these species are involved and that consequently do not occur in the network. The species in 𝒜 and the reactions in ℬ should not be considered in the matrices and vectors describing the network. Therefore, the rows corresponding to 𝒜 and the columns corresponding to ℬ of the stoichiometric matrix ***S*** shall be eliminated, such as the entries of the flux vector ***v*** referred to ℬ. Further, the row referred to the elemental specie must be eliminated both from the moiety vector ***c*** and the conservation laws matrix, whose columns associated to A shall be removed as well.

The Gain of Function (GoF) mutation of a specific species implies that a set of reactions involving it do not occur in the network any longer (Sommariva *and others*, 2021*b*). Let us define with ℬ the set of these reactions. All the chemical species that are involved only in the reactions of ℬ, as either reactants or products, shall be inserted in a set 𝒜 of species to be eliminated from the network. In this case, only the physiological stoichiometric matrix and the fluxes vector are modified from the physiological system. The columns of ***S*** and the entries of ***v*** referred to the reactions in ℬ are eliminated, such as the rows of ***S*** corresponding to the species in 𝒜.

## 3. Results

SSI has been defined in a general setting and therefore can be utilized for sensitivity analysis for all possible CRNs. However, here we focused on applications concerned with the colon-rectal CRN in both healthy and mutated conditions.

### 3.1 Sensitivity analysis for the physiological colon-rectal CRN

We first computed the SSI values in the case of the healthy colon-rectal CRN. As said, in the G1-S phase this network comprises *n* = 419 proteins involved in r=850 chemical reactions and p=81 independent conservation laws. We first pointed out the components of ***k*** and ***c*** characterized by the maximum, median, and minimal SSI values, respectively. The results of this computation are presented in Figure 2 and clearly show that these components are, respectively, the ones with highest, median, and lowest global sensitivity, this latter being assessed as the Euclidean norm of 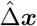.

**Fig. 2.**
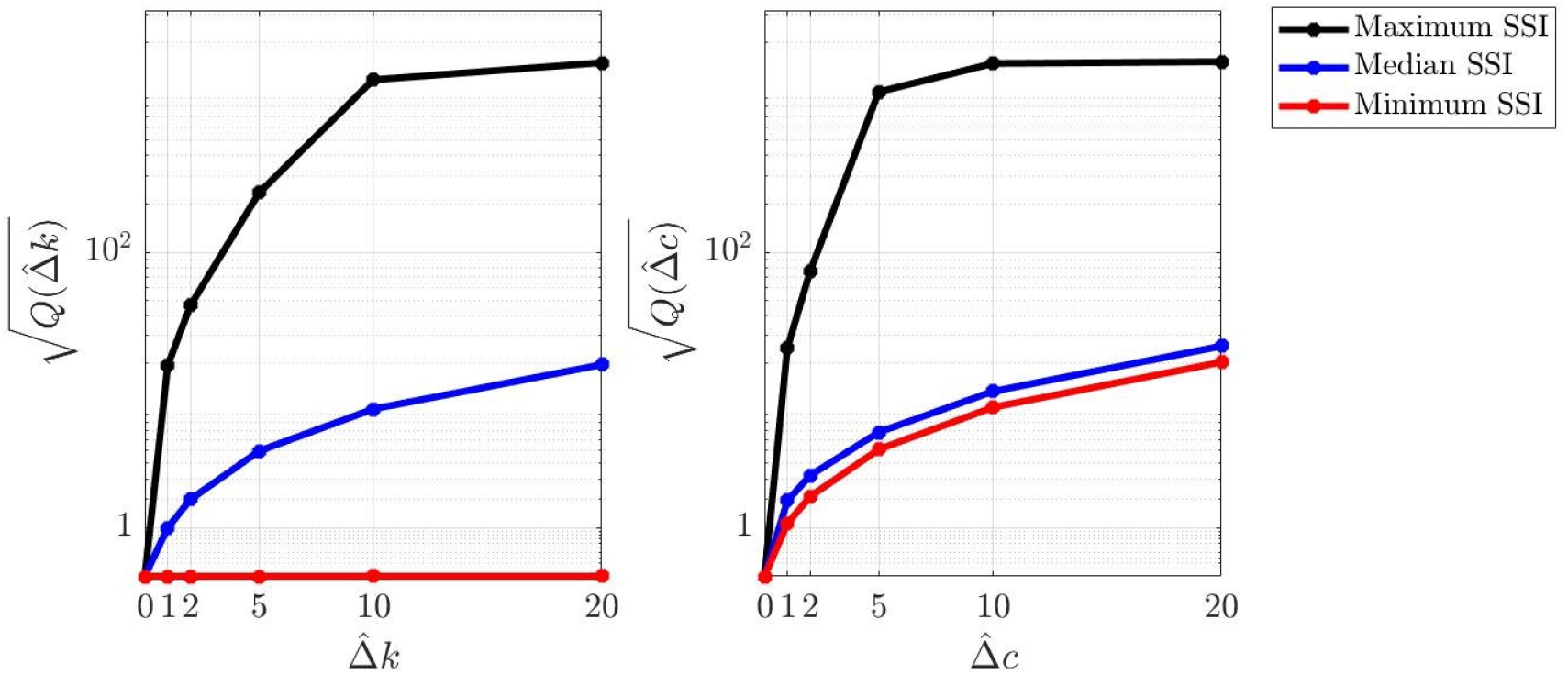
Global sensitivity and SSI values in the physiological colon-rectal CRN. Left panel: 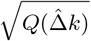 as a function of the relative variation of the rate constants with minimum, median, and maximum SSI values. Right panel: 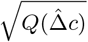 as a function of the relative variation of the conservation laws’ constants with minimum, median, and maximum SSI values.

As said in sub section 2.3.1, one of the nice aspects of SSI is that it allows a reliable estimate of the sensitivity of specific subnetworks of the CRN. For example, Figure 3 shows the results of the same analysis as in Figure 2, but this time focused on the sensitivity of the subnetwork involving gene TP53 (Iacopetta, 2003), which is made by a single equation. Also in this case, the SSI values are nice indicators of the sensitivity level associated to each component of the conservation laws’ vector ***c***. Even more interestingly, in this case the conservation law associated with the maximum SSI value is the one associated to MDM2, which is in fact directly involved in the degradation of TP53 (Sommariva *and others*, 2021*b*). Coherently, a further analysis not shown here demonstrated that TP53 concentration is reduced by increasing the component of ***c*** corresponding to the MDM2 conservation law.

**Fig. 3.**
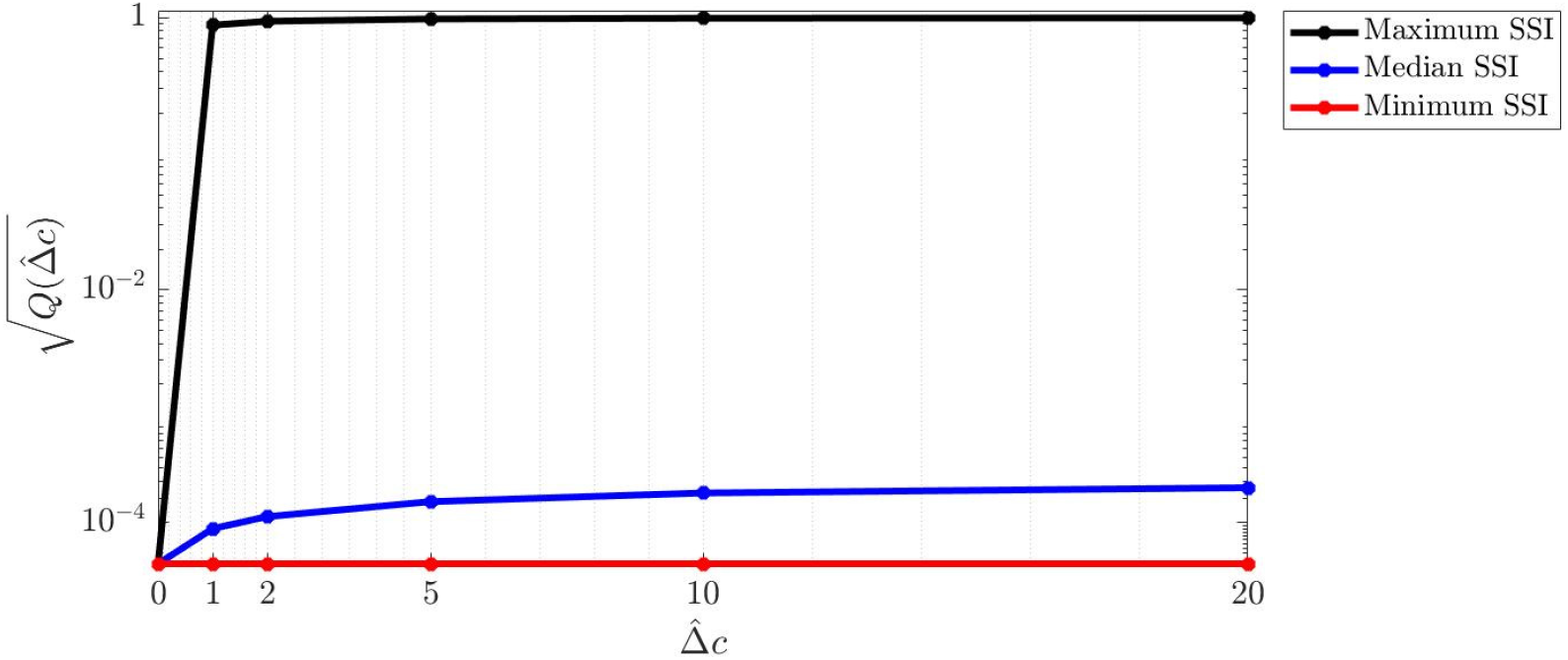
Sensitivity of the equilibrium concentration of TP53: 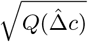 for the TP53 subnetwork is represented versus the relative variation of the conservation laws’ constants with minimum, median, and maximum SSI values.

### 3.2 Sensitivity analysis for the CRC CRN

A greater significance of our approach to sensitivity analysis is concerned with applications to mutated CRN. Indeed, in this context, SSI plays two roles.

As the protein concentration at equilibrium, SSI is an indicator of the impact that each mutation has on the expression of that specific protein. This role is clearly shown by Figure 4 and Figure 5, which represent the SSI values for all components of the rate constant (Figure 4) and conservation laws’ (Figure 5) vectors associated to the colon-rectal CRN when four CRC mutations are included in the network. These figures indicate that the KRAS GoF mutation strongly modifies the SSI values of most indexes and this is coherent with the fact that the concentration values at equilibrium are significantly altered by the presence of such GoF mutation in the network (Sommariva *and others*, 2021*b*).

**Fig. 4.**
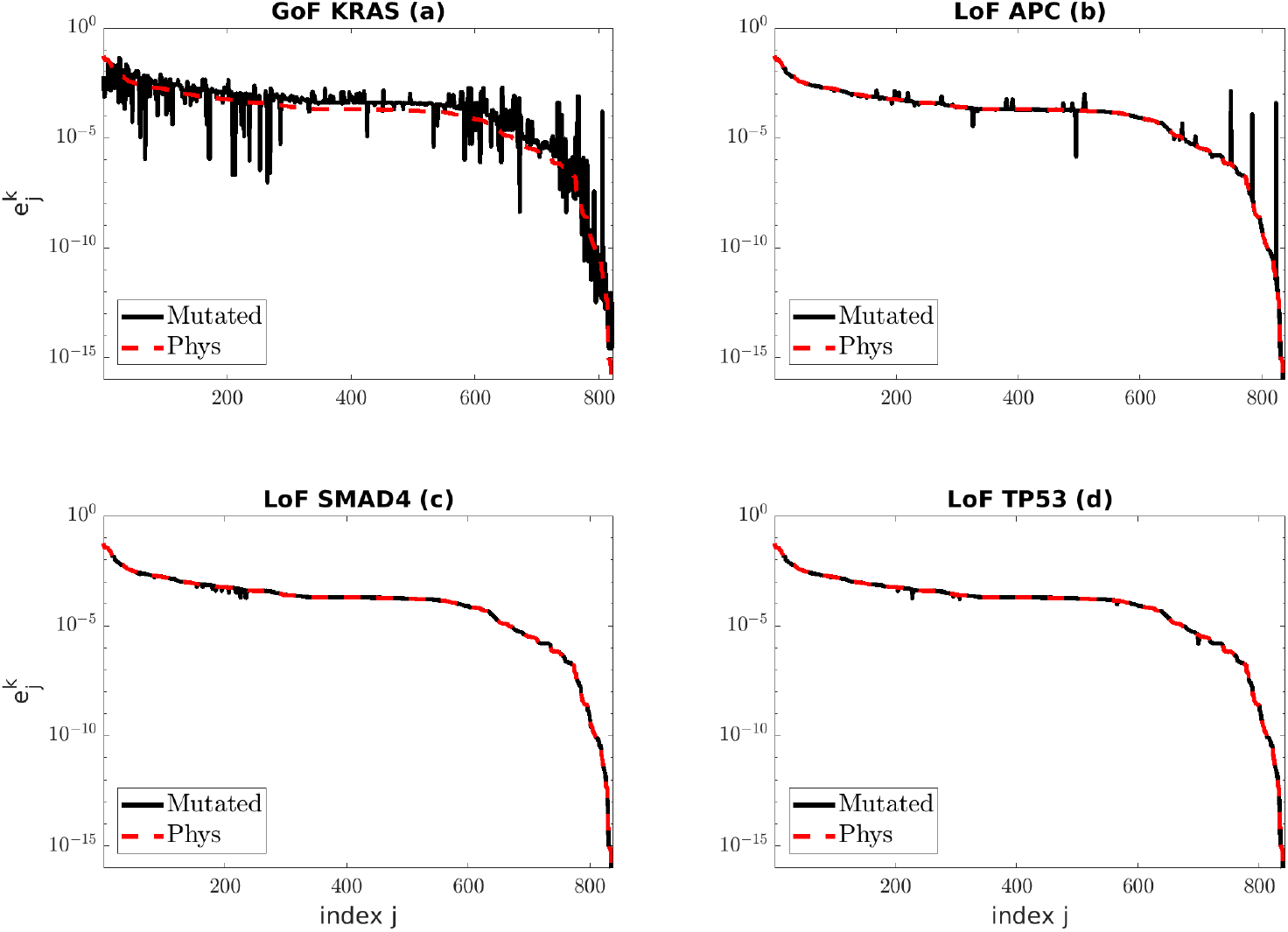
SSI values for all rate constants in the case of four mutated networks (black solid line) with respect to the SSI values of the physiological CRN (red dashed line).

**Fig. 5.**
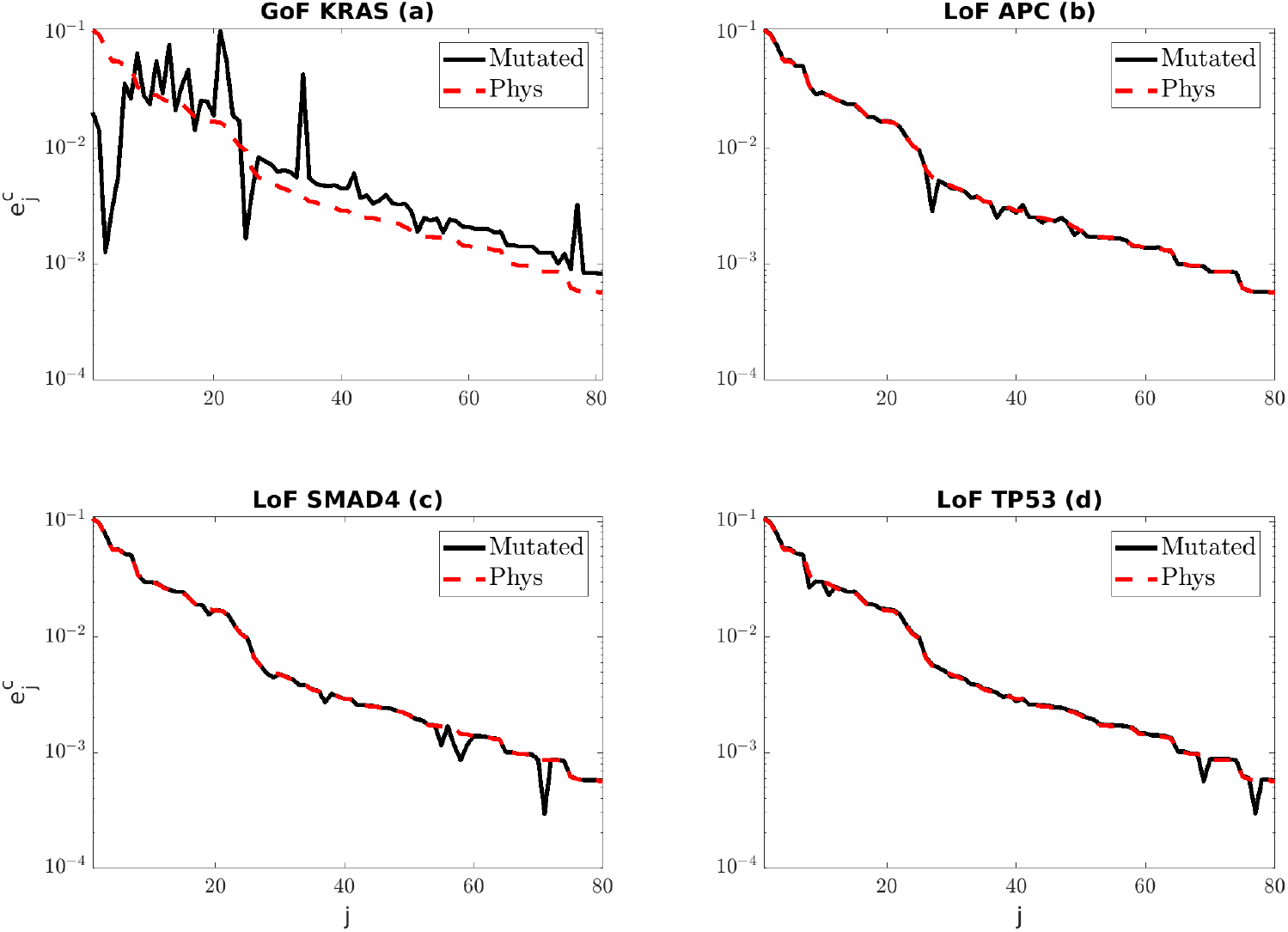
SSI values for all conservation laws’ constants in the case of four mutated networks (black solid line) with respect to the SSI values of the physiological CRN (red dashed line).

More than this, SSI also allows the identification of the specific reactions in the network that were mostly affected by the mutation. This second role of SSI is pointed out by Table 1 that contains a list of reaction ranked according to the values of the relative difference between the SSI values in the physiological and mutated cases (the first column of the table contains the reference numbers attributed to the reactions in the supplementary material of Tortolina *and others* (2015)). All these reactions are related to the signalling pathways of the Ras protein, which is involved in the most important signalling cascades regulating cell growth, proliferation, and survival for many cancer types (Orton *and others*, 2005; Levine *and others*, 2019). Via R41, free Ras is freezed in the inactive chemical compound *Ras*_*GDT*, while R52 leads to a reduction of the active form *Ras*_*GTP*, which is expected to be available in large amounts because of the mutation; the same compound of R52 is generated in reaction R49. Further, among the three reactions R294, R297, and R302, only the last one involves Ras explicitly, via the compound *Ras*_*GDP*. The other two reactions deal with the same compound *ERBP*_*ShP*_*G*_*S* but, more importantly, they are deeply involved in the process of signal transmission from the activation of receptors on the cell membrane to the molecules of Ras. Similar remarks hold true for the remaining reactions R412, R415, R420 (see the Supplementary Material in Tortolina *and others* (2015)).

**Table 1.** Reactions whose sensitivity indexes are mostly effected by the GoF mutation of KRAS. The third column contains the relative differences of 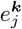 between the mutated and the physiological CRC CRNs.

| Reference # | Reaction | $\delta e_j^k$ |
| --- | --- | --- |
| R41 | Ras + GDP -> Ras_GDP | $1.91e + 07$ |
| R294 | ERBP_ShP_G_S -> ERBP_ShP_G + SOS | $2.91e + 04$ |
| R49 | RP_ShP_G_S_Ras + GTP -> RP_ShP_G_S_Ras_GTP | $2.88e + 04$ |
| R412 | ERB3P_Sh -> ERB3P + Shc | $2.86e + 04$ |
| R415 | ShP + ERB3P -> ERB3P_ShP | $1.23e + 04$ |
| R297 | ERBP_ShP + G_S -> ERBP_ShP_G_S | $8.64e + 03$ |
| R302 | ERBP_ShP_G_S_Ras + GDP -> ERBP_ShP_G_S_Ras_GDP | $5.78e + 02$ |
| R57 | RP_G_S_Ras_GDP -> RP_G_S_Ras + GDP | $5.78e + 02$ |
| R420 | ERB3P_ShP_G + SOS -> ERB3P_ShP_G_S | $5.78e + 02$ |
| R52 | RP_ShP_G_S + Ras_GTP -> RP_ShP_G_S_Ras_GTP | $4.95e + 02$ |

## 4. Discussion and conclusions

CRNs are typically characterized by a huge number of variables, and although static formulations reduce the number of parameters in the game, yet calibration of a CRN is often a challenging issue. Sensitivity analysis of both physiological and mutated CRNs is an effective way to reduce the complexity of the network, since just sensitive rate and conservation laws’ constants should be regarded as parameters to finely tune, while insensitive variables can be coarsely approximated.

In our view, the sensitivity index introduced in this paper has several advantages. It is specifically tailored to the static case and it is based on PCA, which makes its computation rather straightforward. From a technical viewpoint, it does not suffer any intrinsic sign ambiguity, as it occurs in the case of the index introduced by Liu *and others* (2005). Further, it can be applied for the sensitivity analysis of a specific single species, of a specific pathway, and of the whole network; and, also, it can be used to nicely rank the reactions with respect to the impact that a specific mutation has on the signalling pathways. Finally, although we have validated SSI in the case of the CRC CRN, this index has a more general value, and can be utilized for assessing the equilibrium condition of any mutated signalling network.

As far as future work is concerned, the next step will be to use SSI to select the sensitive pathways within the global CRC CRN, and then to calibrate the simplified network by applying computational algorithms based on inverse problems theory (Bertero and Piana, 2006) to experimental measurements of the proteins’ concentrations at equilibrium.

## Acknowledgements

The support of the Ecosistema ‘RAISE – Robotics and AI for Socio-economic Empowerment’ (PNRR - Missione 4 Componente 2 Investimento 1.5) is kindly acknowledged. S.S. and M.P. have been partially supported by Gruppo Nazionale per il Calcolo Scientifico (GNCS-INdAM)

## Software

Software in the form of Matlab code is available at https://github.com/theMIDAgroup/CRN_sensitivity

